# Spatial and temporal variation in small mammal abundance and diversity under protection, pastoralism and agriculture in the Serengeti Ecosystem, Tanzania

**DOI:** 10.1101/727206

**Authors:** Monica T. Shilereyo, Flora J. Magige, Joseph O. Ogutu, Eivin Røskaft

## Abstract

Land use is an important factor influencing animal abundance, species richness and diversity in both protected and human-dominated landscapes. Increase in human population and activities intensify changes in habitat structure and hence abundance, species richness and diversity. We investigated the influences of land use and seasonality on small mammal abundance, species richness and diversity in 10 habitat types distributed over protected, agricultural and pastoral landscapes in the Serengeti ecosystem in Tanzania. We used live traps (*n* = 141) and capture-recapture methods in each of 10 fixed plots distributed across three landscapes for a total of 28,200 trap nights of effort. Trapping was carried out in the wet and dry seasons for two consecutive years (April 2017 to October 2018). Small mammal abundance was higher in the pastoral than in the protected and in the agricultural landscape. Abundance was higher in the dry than the wet season across all the three landscapes. Species richness and diversity were higher in the protected, middling in the agricultural and lowest in the pastoral landscape. The high abundance in the pastoral landscape was due to the numerical dominance of two species, namely A. *niloticus* in the shrubland and *M. natalensis* in the cropland habitat, resulting in low species richness and diversity. Abundance was more evenly distributed across all habitats in the protected area due to less disturbance. The low abundance in the agricultural landscape, likely reflects disturbance from cultivation. High species richness and diversity in the protected area indicate high habitat heterogeneity while high species diversity in the agricultural landscape was likely due to high food availability during and soon after harvests. These findings emphasize the importance of protection in maintaining habitat heterogeneity for wildlife. They also reaffirm the need for buffer zones around protected areas to cushion them from intensifying human activities.

## Introduction

Human influence on ecosystems is increasing worldwide due to rapid population growth and increasingly resource-consuming life styles (1). This influence has become so important that mankind is now and, will likely remain for years to come, the main global driver of ecological change (2, 3). Human-altered ecosystems made of various settlements, agro-pastoral and protected areas dominate the terrestrial biosphere, covering more than three quarters of the total ice-free land areas (4). These alterations to ecosystems have resulted in a global biodiversity crisis that threatens the world’s species and ecosystems (5-7). Today, most protected areas, set aside to safeguard the remaining global biodiversity, are surrounded by different human activities making them isolated “islands”. This change raises fundamental questions concerning whether all protected areas will last into the far future given the current rate of increase in human population and activities (8-12).

Human activities that cause land use change also act as drivers of biodiversity loss (13). Agriculture is the dominant land-use activity on the planet and is responsible for altering and endangering wildlife communities on a massive scale (14, 15). It has transformed native vegetation into monocultures thereby decreasing biodiversity by homogenising habitats (16).

Although, agricultural activities can provide food to some wildlife species, a leading conservation concern is that agricultural lands alter wildlife communities, favouring generalists at the expense of specialists (17, 18). On the other hand, livestock grazing, apart from promoting vegetation regrowth and nutrient enhancement, causes mechanical disturbance, reduces plant biomass and changes vegetation composition (19). The changes in vegetation structure can have several knock-on effects on critical ecosystem functions, such as provision of shelter and food for wild animals, species composition and richness (20-22).

Small mammals have long been used as bioindicators and model organisms to study patterns of species abundance and diversity along different land use gradients (23-26). These studies show that both grazing and farming activities differentially influence small mammal community characteristics, such as species richness, diversity and abundance (14, 18, 27-31). In particular, in the Serengeti National Park in Tanzania, small mammal studies have focussed on species and biotope, diversity and abundance in different habitats and along altitudinal gradients (32, 33); human-small mammal conflicts (34) and influence of small mammals on their predator abundances (35). A few studies have also compared protected areas with their adjacent human-dominated habitats to infer the influence of anthropogenic activities on small mammal species diversity and abundance (17, 27, 32). Assessments of the influence of human activities on small mammal species diversity, richness and abundance have produced mixed results, ranging from positive, negative to neutral effects. This is unsurprising given the complex and dynamic interactions among ecological, historical, and evolutionary processes shaping rodent diversity (36).

Surprisingly, few studies have sampled small mammals simultaneously between protected areas and the adjoining human-inhabited areas across seasons (17, 32). This study aims at expanding upon the earlier studies by assessing spatial and temporal variation in small mammal species diversity, richness and abundance in the protected and adjoining human-dominated livestock grazing and agricultural landscapes in the Serengeti ecosystem. We address the following two objectives. First, we quantify the species richness, diversity, abundance and composition of small mammals in 10 habitats distributed across the three land use types. Second, we analyse temporal variation (seasonal and interannual) in small mammal abundance and diversity across the 10 habitats and three land use types. We anticipate that if disturbance reduces structural and functional habitat heterogeneity then small mammal species diversity and population density should be highest inside the protected areas, intermediate in the pastoral lands and lowest in the cultivated areas. In addition, since small mammals exhibit pronounced reproductive seasonality such that more juveniles are produced during the early dry season (June and July) we expect to find a higher density of most of the species in the dry than the wet season because of elevated food abundance linked to higher rainfall in the wet season. Nevertheless, we anticipate that species should respond to human disturbance in contrasting ways, such that habitat generalists should be able to colonize disturbed areas faster than habitat specialists. Thus, we expect the abundance of habitat generalists to be higher than those of specialists in the more disturbed pastoral and cultivated lands than the protected land.

## Materials and Methods

### Study area

Data were collected in the Greater Serengeti Ecosystem in Tanzania, East Africa. Our focus was on the north-eastern Serengeti ecosystem; including the Serengeti National Park (2° 20′ S, 34° 50′ E) and two adjacent administrative districts, namely the Serengeti (2°15′ S, 34°68′ E) and Ngorongoro (3°24′ S, 35° 48′ E). Serengeti National Park protects 14750 km^2^ of tropical savanna ecosystem (37). The park comprises woodlands and open grasslands, besides other more restricted habitat types (35, 38), with farming and livestock herding practiced around the ecosystem.

The study covered mainly the northern part of the Serengeti ecosystem within three main blocks located along the Mto wa Mbu-Musoma road. This area was selected because it contains contrasting land use types, including agricultural areas (south west), pastoral and limited agricultural areas in the south east and the Serengeti National Park situated in-between these two blocks (Fig. 1). The Mto wa Mbu-Musoma road bisects each of the three blocks, resulting in 6 sub-blocks; three sub-blocks on either side of the road. Based on habitat type, we selected two study plots from each of the 6 sub-blocks resulting in 12 study plots. However, only 10 plots were included in the study because the other two plots (wooded grassland and grassland), situated in Ololosokwan; a pastoralist village with a historical land use conflict with the Tanzania National Parks, were excluded.

**Fig 1.**
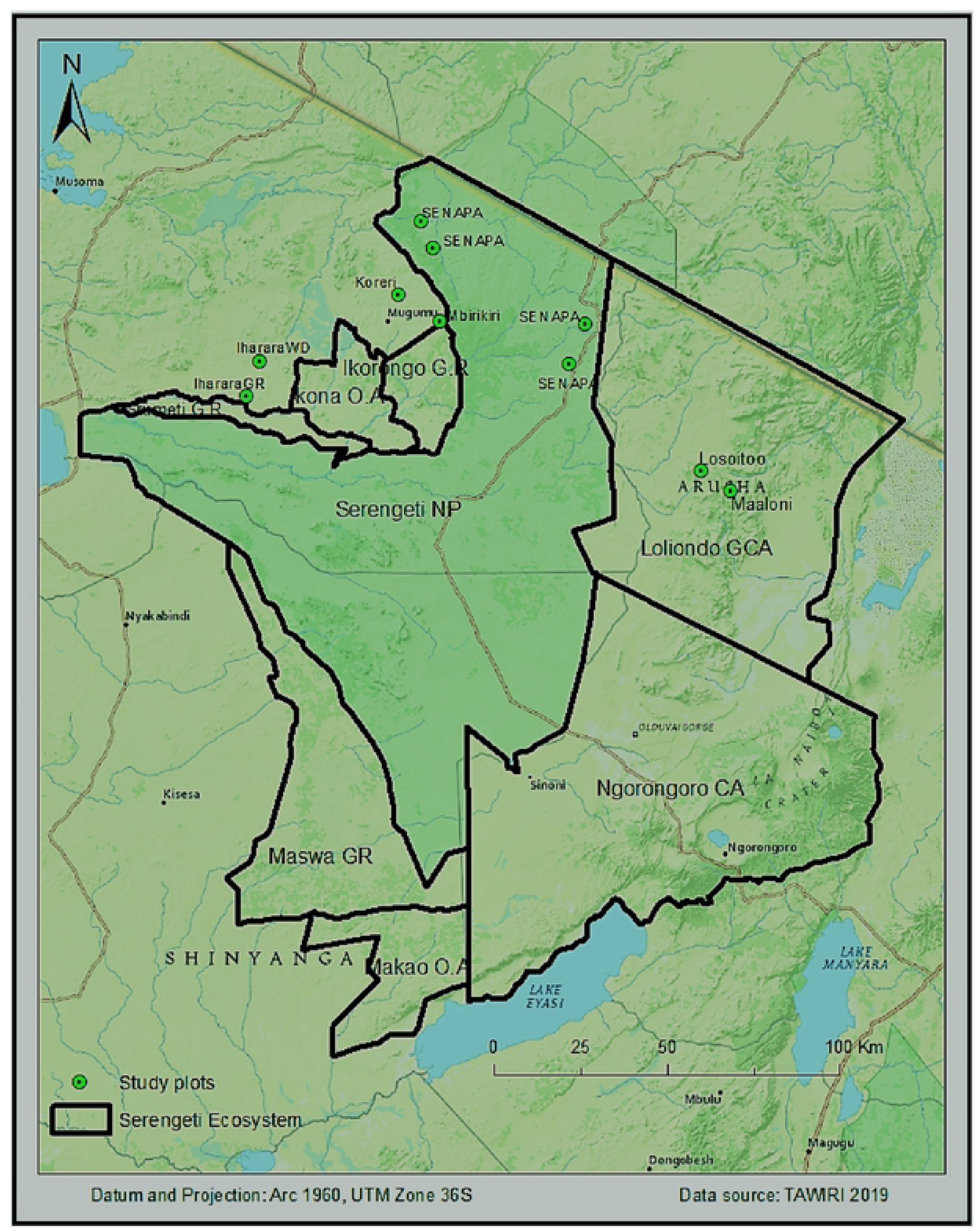
Map of the Serengeti Ecosystem showing the study area.

Green circles with black dots inside indicate the study plots inside and outside the Serengeti ecosystem

The climate in the ecosystem is warm and dry, with mean annual temperatures varying between 15 °C and 25 °C (27). The rainy season is bimodal with the short rains spanning November - January and the long rains covering March - May (39). Rainfall increases from east to west towards Lake Victoria south to north and south-east to north-west. Rainfall increases along a south east-north west gradient from 800 mm / year on the dry south-eastern plains to the wet north-western section (1,050 mm / year) of the Serengeti National Park (40). However, during this study (April-May and August-September 2017 and 2018) the mean monthly rainfall averaged 153 mm while the temperature averaged 26 °C.

### Ethical clearance

The study design was approved by Tanzania Wildlife Research Institute (TAWIRI) and the permit to conduct research was obtained from Tanzania National Parks (TANAPA). For the private land, the permit was issued by District Executive Office: Serengeti and Ngorongoro Districts. All captured small mammals were handled according to the approved permit and released immediately at the point of capture after observation.

## Methods

### Trapping procedures

Each land use consisted of 4 plots (except the pastoral landscape that had only 2 plots), each measuring 100 × 100 m and selected in representative habitat types, including grassland, shrubland, wooded grassland, cropland and riverine forest habitats. Except for the pastoral land use where only two habitats were sampled (cropland and shrubland), other land use types had four habitats each; wooded grassland, grassland, cropland and shrubland in the agricultural and wooded grassland, grassland, riverine forest and shrubland in the national park. A total of 141 small mammal traps (100 Sherman traps, 30 wire mesh traps and 11 bucket pitfall traps) were set in each of the 10 plots for five consecutive nights and then transferred to the next plot. Trapping was done twice a year, April-May for the wet season and August and September for the dry season, for two consecutive years (2017 and 2018). We started trapping on 18^th^ April 2017 and stopped on 20^th^ September 2018. Trapping started on the pastoral landscape (eastern part of the Serengeti ecosystem) followed by the protected area and then by the cultivated landscape (western part of the ecosystem) because the eastern part of the ecosystem receives relatively low rainfall and so gets drier early compared to the western part. The same pattern was followed except for one season (wet season 2018) due to logistical constraints, which forced us to set traps in the protected area after the agricultural landscape.

Pitfall lines and trap lines were installed to capture mostly shrews and rodents, respectively. Each plot was assigned one pitfall line consisting of 11 buckets, placed 5 m apart, and buried in the ground so that the top of the bucket was at the ground level. Each of the 22 buckets per plot was 26 cm deep and had upper and lower diameters of 30 cm and 26 cm, respectively, and a 20-litre capacity. The bottom of buckets was pierced with small holes to allow water drainage. Each pitfall line had a 50 cm-high black plastic drift fence running over the center of each bucket. These passive and non-baited traps capture animals moving on the habitat floor that encounter the drift fence and follow it until they fall into a bucket. The pitfall lines were generally set along straight trails; however, rocks and logs occasionally forced deviations. This technique has been used with considerable success in other small mammal surveys (41, 42). For the Sherman traps (23 × 9.5 × 8 cm), 10 lines (10 m apart) were developed on the grid. Sherman traps were arranged along the lines, with a total of 100 traps placed on a 100 × 100 m plot and spaced 10 m a part. To maximize capture and variety of small mammals caught, 30 wire mesh traps ‘*Mgono*’ were placed in-between the Sherman trap lines. Five wire mesh traps were placed 20 m apart from each other. These wire mesh traps are widely used in Tanzania by local hunters, and are funnel-shaped, multi-capture traps made of thin wire. Bait for both the Sherman and ‘*Mgono’* traps consisted of freshly fried coconut coated with peanut butter and mixed with sardines. Traps were rebaited every morning and evening.

Checking of traps was done twice a day, early in the morning and evening. Equal amounts of time were allocated to both methods, so we use ‘trap-night’ (one trap in operation for one 24-hr period, 0700 to 0700 hrs, to quantify sampling effort). We refer to the success rate of capture as trap success and calculate it by dividing the number of individuals captured by the number of trap-nights and multiplying by 100. Trap success has been recommended as a good measure of spatial and temporal variation in relative abundance (43). Traps stayed in one plot for 5 consecutive days before being taken to the next plot. Using recorded morphometric (external shape and dimensions) measurements and field guides we identified trapped animals to genus or species (44). In addition, distinguishing features like species, sex, size, reproductive status, and presence of scars or particular characteristics were recorded to facilitate individual identification (45). We marked trapped animals by toe clipping and released them at the points of capture. After a standardized procedure, involving live trapping and a complete dataset of small mammal abundance in each land use type, we aimed to ascertain the influence of human activities on species richness, diversity and abundance.

### Statistical analyses

To establish the pattern of small mammal response to abiotic and biotic factors, we analysed variation in abundance, species diversity and richness across the three land use types, 10 habitat types and two seasons. Captures from the same land use and habitat type were pooled together and represented by the frequency for the particular land use or habitat type. Data were analysed using *R* version 3.5.2. Reshape and Dplyr packages (46) were used to calculate descriptive statistics whereas the iNEXT package was used to calculate species diversity and richness; Chao richness order 0,1 and 2, among the land use types and habitats (47). The method efficiently uses all the available data to make robust and meaningful comparisons of species richness between assemblages for a wide range of sample sizes or completeness. Also, it has been generalized to diversity measures that incorporate species abundances and those that take into account the evolutionary history among species (48). Hutcheson-t test was used to test the significance of differences in diversity across the three-land use types and habitat types.

Chi-square goodness-of-fit tests were used to test whether the observed abundances differed significantly from expectation assuming a uniform distribution. Chi-square tests were followed by the chisq.multcomp post hoc test from the RVAideMemoire package (49). Abundance in each of the three land use types were corrected for differences in trapping efforts and the results presented as the number of small mammals/ 100 trap nights. In this study, the significance level of 0.05 was adopted.

## Results

### Small mammal species richness and diversity

Overall, the species richness (*S*) was 19 species, of which 15 species were recorded in the NP, 11 in the AG and only 9 in the PA, but the difference in species richness among the landscapes was not statistically significant 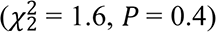. The overall diversity (H’) was 2.25 and varied across the three landscapes (Fig. 2). Species diversity was twice as high both in the NP (Hutcheson’s t-test, *t*_513_ = 8.0, *P* < 0.001) and in the AG (*t*_332_= 7.0, *P* < 0.001) than in the PA landscape but was similar between the NP and AG landscapes (*t*_290_ = 1.2, *P* = 0.22). In addition, evenness was high in both the AG (85%) and NP (60%) landscapes but low in the PA (30%) landscape an indication of lower dominance in the NP and AG (S1Table).

**Fig 2.**
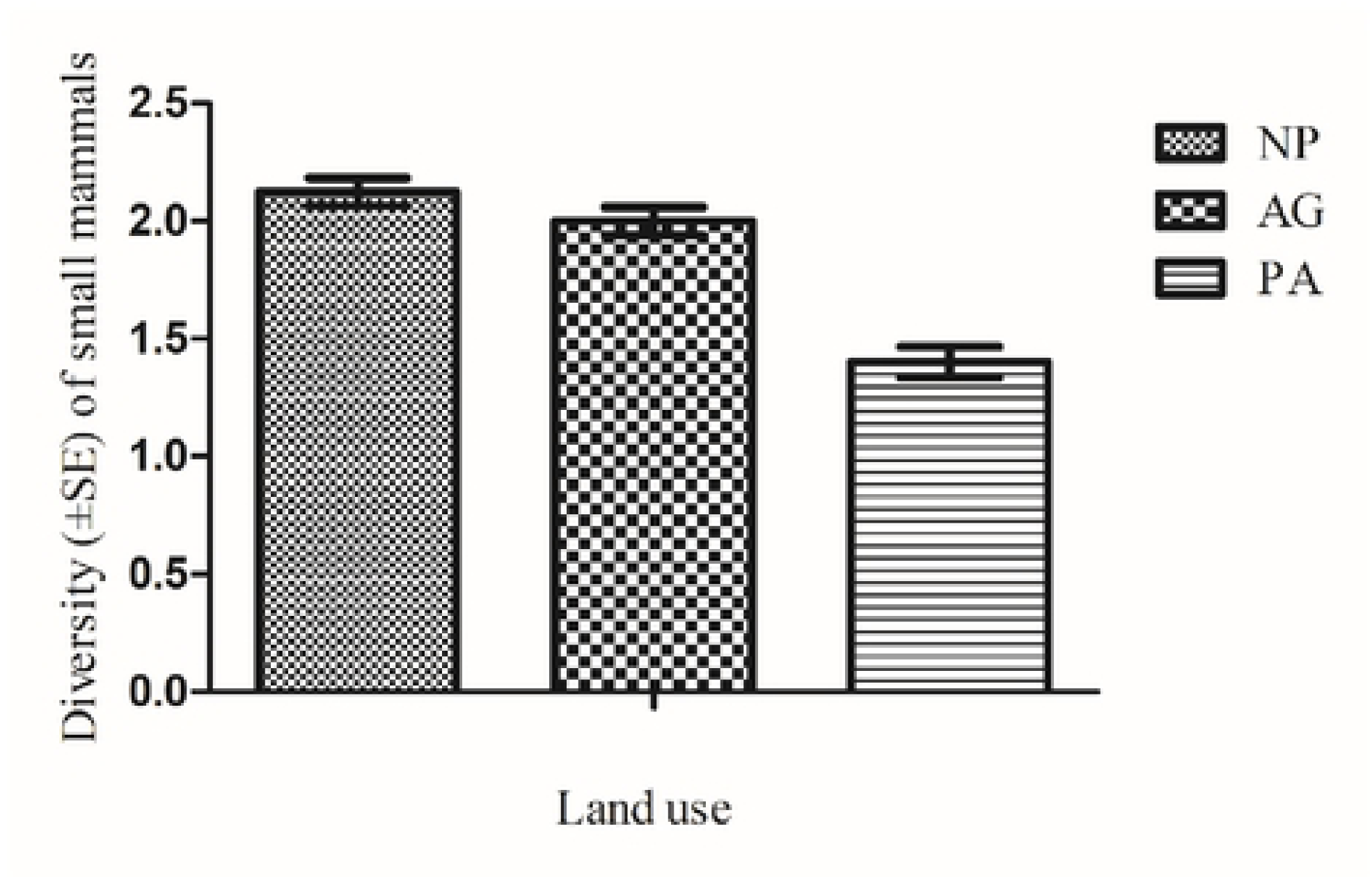
Diversity index (± standard error) for small mammals in three landscapes in the Serengeti ecosystem. AG = Agriculture, NP = National park and PA = Pastoral land use types.

Species richness and diversity also varied noticeably across different habitats in the same land use type and across the same habitat in different land use types. Specifically, in the NP species richness was the highest in the wooded grassland followed by the forest, grassland and shrubland habitats, in decreasing order (Fig. 3a & 3b). However, these apparent differences in species richness were statistically insignificant 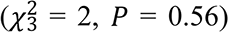. In contrast to richness, species diversity was the highest in the forest, and the lowest in the shrubland habitat. Species diversity was lower in the shrubland than in the forest (*t*_332_ = 5.6, *P* < 0.001), wooded grassland (*t*_129_ = 2.66, *P* = 0.0086), and the grassland (*t*_39_ = −2.14, *P* = 0.038) habitats but comparable between forest and wooded grassland, (*t*_121_ = −1.78, *P* = 0.07) and between forest and grassland (*t*_37_= −1.4, *P* = 0.16) habitats (Table A1).

**Fig 3a.**
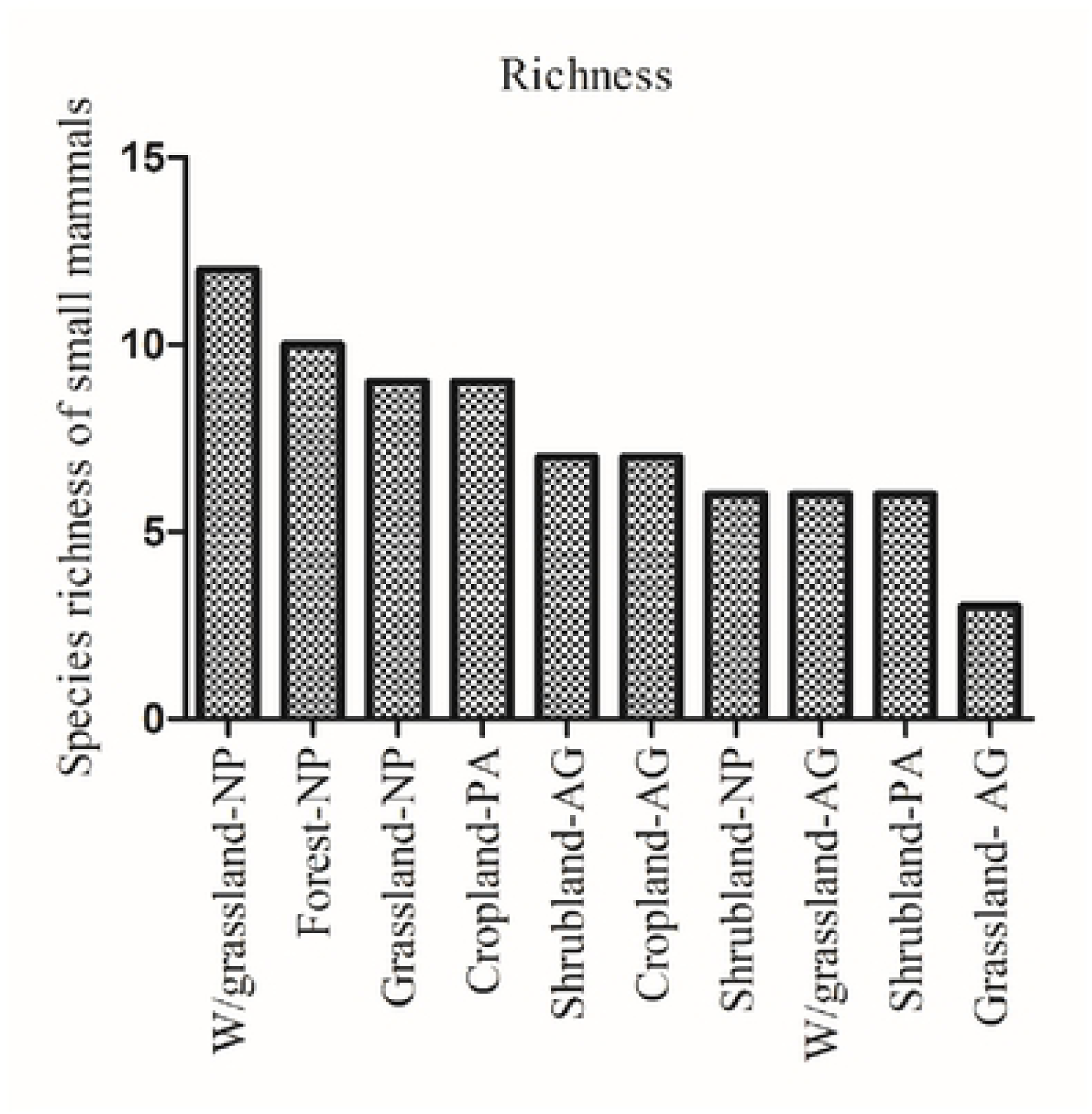
Small mammal species richness in the 10 different habitats across the three land use types. AG = Agriculture, NP = National Park and PA Pastoral landscapes.

**Fig 3b.**
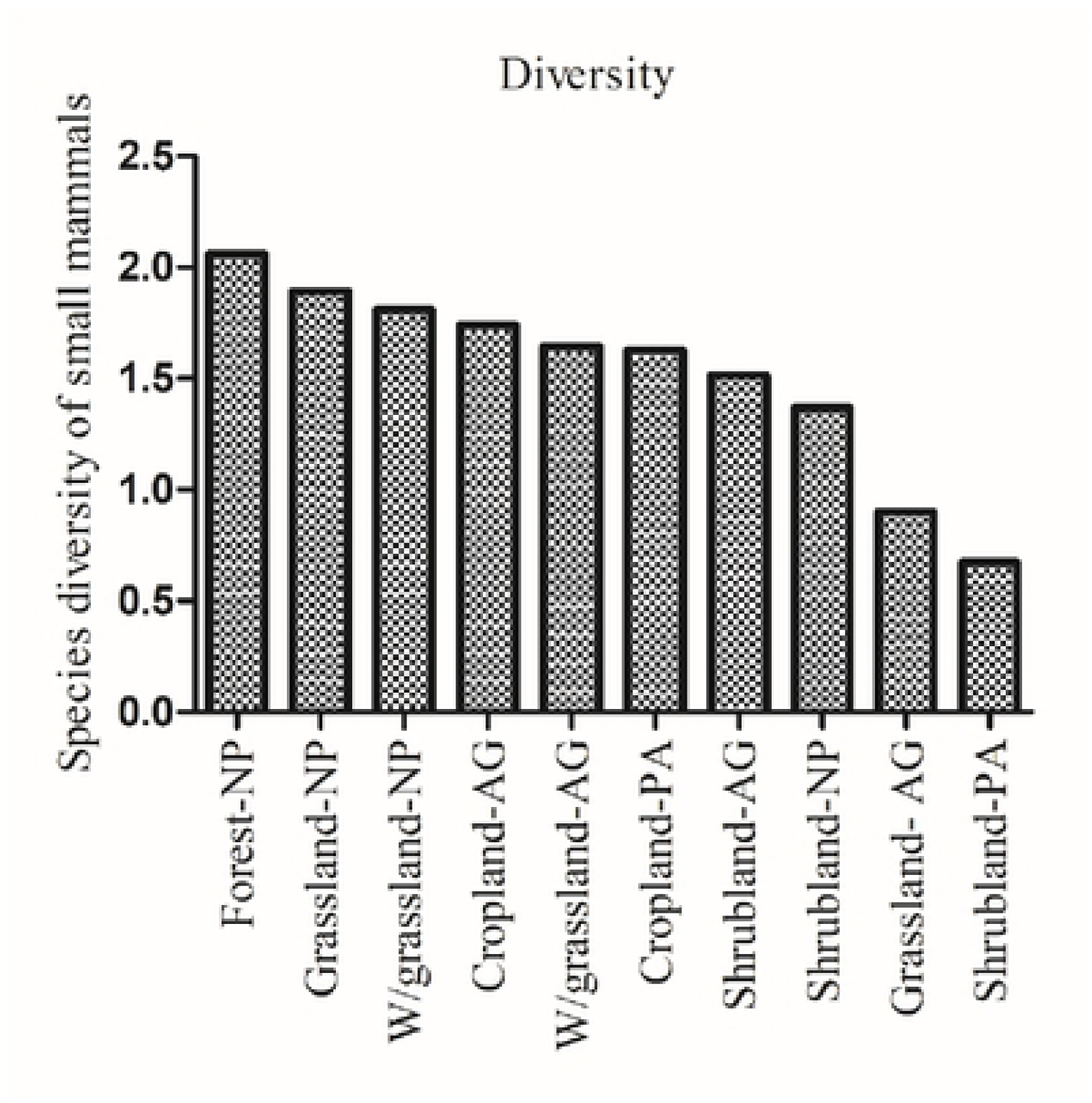
Small mammal diversity in the 10 different habitats across the three land use types. AG = Agriculture, NP = National Park and PA Pastoral landscapes.

For the AG land use type, the wooded grassland (*S* = 6) and grassland (*S* = 3) habitats each had half the number of species found in the same habitat in the NP landscape. Species diversity was significantly lower in the grassland than in the cropland (Hutcheson *t*-test, *t*_17_ = 3.5, *P* = 0.0023), shrubland (*t*_21_ = −2.5, *P* = 0.019) or wooded grassland (*t*_18_= 2.9, *P* = 0.008) habitat. In the PA, species richness was comparable between the shrubland and cropland habitats 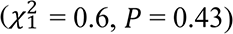. But species diversity was lower in the shrubland habitat in the PA than in the same habitat in the NP (*t*_185_ = −5.7, *P* < 0.001) or the AG (*t*_63_ = −4.6, *P* < 0.001) landscape (Fig. 3b). In addition, species diversity was similar in the cropland habitat in the PA and AG landscapes and other habitats in the NP except for the riverine-forest habitat, which had the highest recorded diversity.

### Small mammal abundance and species composition

An aggregate sampling effort of 28,200 trap-nights spread across all the three land use types resulted in the trapping of a total of 612 individuals belonging to 19 species and two orders (Rodentia and Eulipotyphla) of small mammals. Of these, 86% (n = 528) were rodents whereas 14% (n = 84) were shrews. The number of small mammals captured/100 trap nights was the highest for the pastoral landscape (PA) (4, *n* = 277; trap success = 4.91), followed by the national park (NP) (2, *n* = 237; trap success = 2.2) and the lowest for the agricultural landscape (AG) (0.8, *n* = 98; trap success = 0.84, Table 1).

**Table 1.** Trap success (100 × number of captures/numbers of trap nights) for each species of small mammal recorded for each of the three land use types in the Serengeti ecosystem during the wet and dry seasons of 2017 and 2018. AG and NP had 11,280 trap nights each whereas PA had 5640 trap nights. A “trap night” is one trap set for one full day.

| Species | Trap success |  |  |
| --- | --- | --- | --- |
|  | ‡AG | NP | PA |
| *Aethomys sp* | 0.05 | 0.04 | 0.41 |
| *Arvicanthis niloticus* | 0 | 0.27 | 2.34 |
| *Crocidura sp* | 0.1 | 0.6 | 0.04 |
| *Dendromus melanotis* | 0.06 | 0.35 | 0 |
| *Gerbilliscus vicinus* | 0.195 | 0 | 0.14 |
| *Grammomys sp* | 0 | 0.06 | 0 |
| *Graphiurus murinus* | 0 | 0.15 | 0.05 |
| *Lemniscomys striatus* | 0 | 0.05 | 0 |
| *Mastomys natalensis* | 0.06 | 0.05 | 1.56 |
| *Mus sorella* | 0.13 | 0.14 | 0.07 |
| *Mus sp* | 0.15 | 0.31 | 0.12 |
| *Myomys sp* | 0.07 | 0.01 | 0 |
| *Otomys angoniensis* | 0 | 0 | 0.14 |
| *Praomys jacksoni* | 0 | 0.1 | 0.02 |
| *Rattus rattus* | 0.02 | 0 | 0 |
| *Saccostomus sp* | 0 | 0.05 | 0 |
| *Steatomy parvus* | 0 | 0.01 | 0 |
| *Zelomys sp* | 0 | 0.01 | 0 |
| *Zelotomys sp* | 0 | 0 | 0.02 |
| Total | **0.835** | **2.2** | **4.91** |

‡AG = Agriculture, NP = Serengeti National Park and PA = Pastoral Landscape.

Thus, the most common species had the highest trap success in the pastoral land use and the lowest in the agricultural areas, with a few exceptions. Notably, the trap successes for *M. natalensis* and *A. niloticus* were the highest for both the pastoral and agricultural landscapes whereas those for *Crocidura sp, D. melanotis* and *G. murinus* were the highest for NP. In addition, rare species represented by a total of less than 10 captured individuals, were mostly restricted to within the confines of the protected area, indicating greater diversity. Specifically, the rare species were *Grammomys sp, Lemniscomys striatus, Praomys jacksoni, Saccostomys sp* and *Zelomys sp.*

Trap success differed significantly across the three land use types (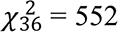, *P* < 0.001, Fig 4). In particular, it was lower for the AG than for either the PA 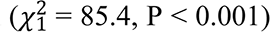, or the NP landscape 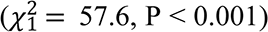 and higher for the PA than the NP land use 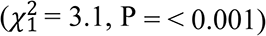. Hence the abundance of small mammals decreased from the PA through the NP to the AG landscape. Note that even though the PA had the highest number of small mammals, it is species poor because only two species (*A*. *nilotocus* and *M. natalensis*) made the most contribution (80%) to the total capture. This contrasts with the NP landscape where the most common species (*D. melanotis* and *Crocidura spp*) contributed only 44% to the total capture, meaning a more even contribution of species to the overall abundance (S2Table). However, when only either or both species, *A. niloticus* and *M. natalensis*, were excluded from the analysis, NP had a significantly higher abundance of small mammals than the other two land use types. When both species were omitted from analysis, the abundance of small mammals was still significantly different between the land use types 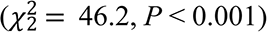, but became higher for the NP than the AG 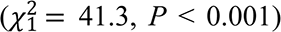 or the PA 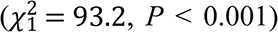 landscape. Therefore, the pattern of abundance changed such that abundance became the highest for the NP, middling for the AG and lowest for the PA landscape.

The abundance of small mammals also varied between different habitats within each land use type, but the pattern of the differences was inconsistent across the three land use types. For the NP landscape, the abundance of small mammals varied across habitats 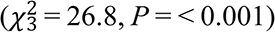 such that it was lower in the grassland than in the wooded grassland, shrubland and forest habitats (Fig. 5). For the AG, the abundance of small mammals differed significantly across the four habitats 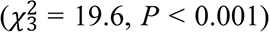 and was higher for the shrubland than for the other habitats (Fig. 6). However, there was no difference in the abundance of small mammals among the wooded grassland, cropland and grassland habitats or between the cropland and shrubland habitats 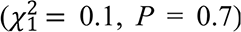. The latter two habitats had the highest abundance of small mammals in the ecosystem, dominated by *A*. *niloticus*, which contributed 78% of all the captures in the shrubland and 46% of *M. natalensis* in the cropland (S2Table).

**Fig 4.**
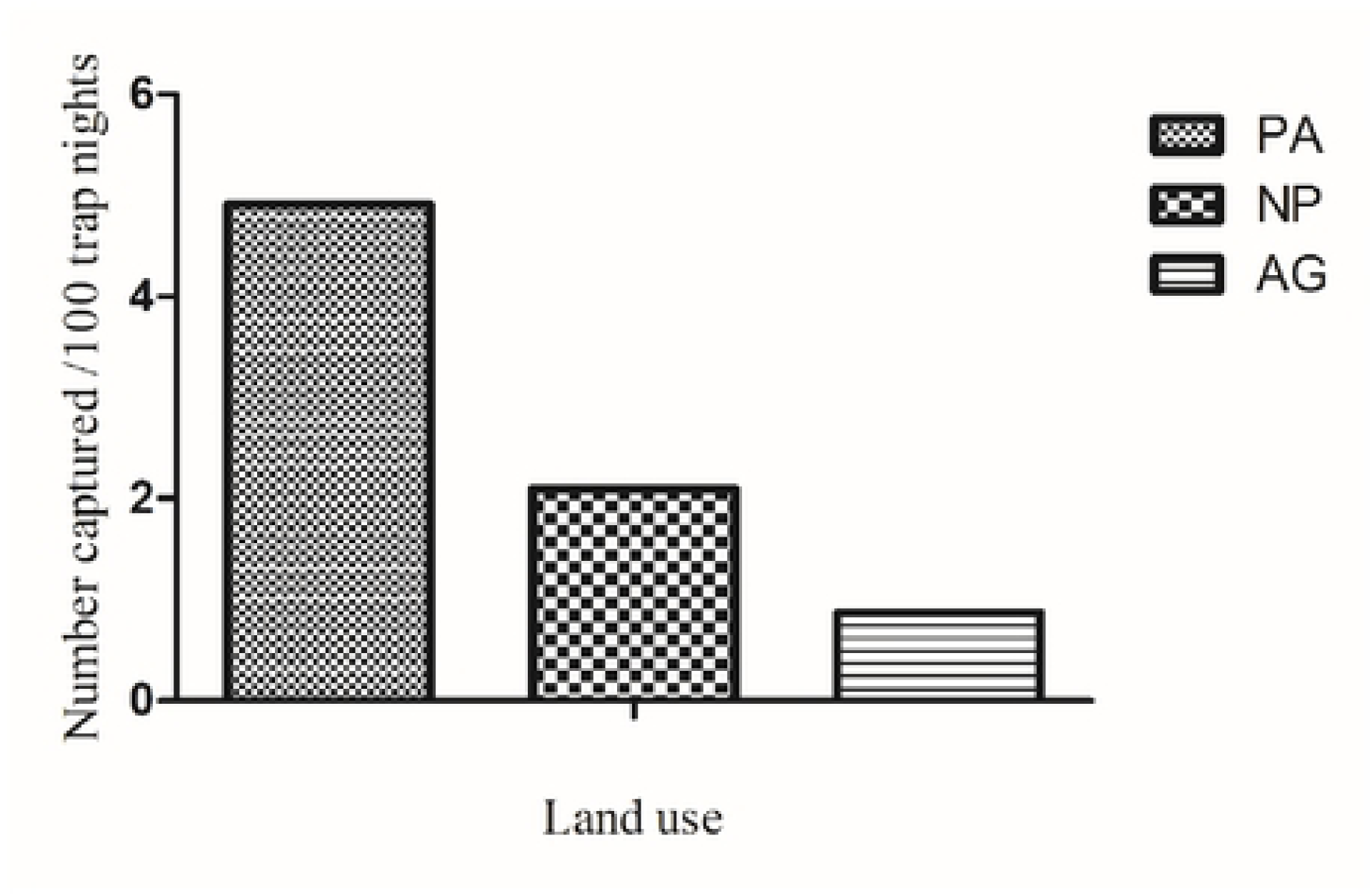
Number of small mammals caught per 100 trap nights in each of the three land use types. AG = Agriculture, NP = National Park and PA = Pastoral landscape.

**Fig 5.**
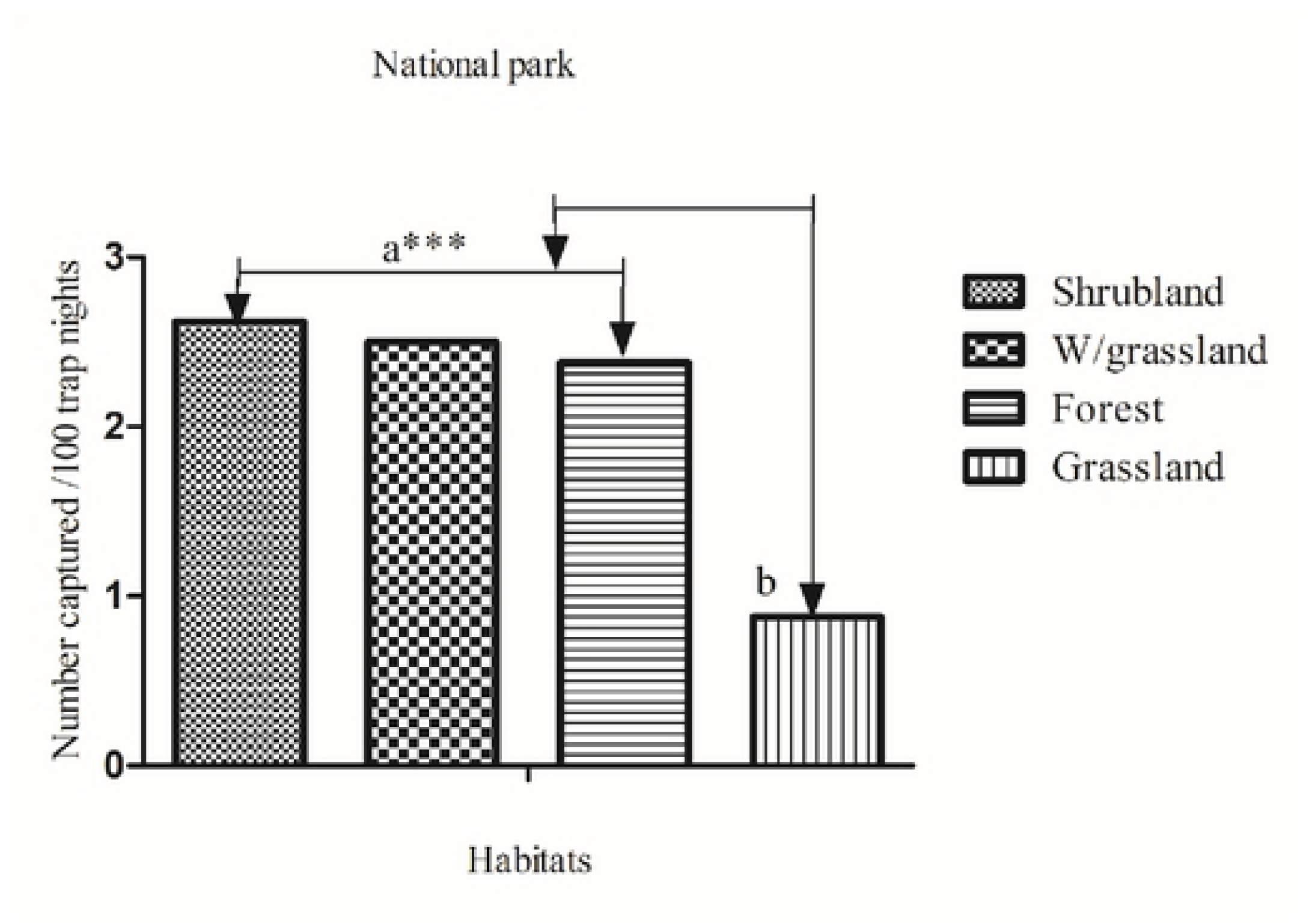
Number of small mammals caught per 100 trap nights (abundance) in each habitat in the NP landscape. Grassland (a) is statistically significantly different from all the other three habitats (b) (*** indicates level of significance (P < 0.001).

**Fig 6.**
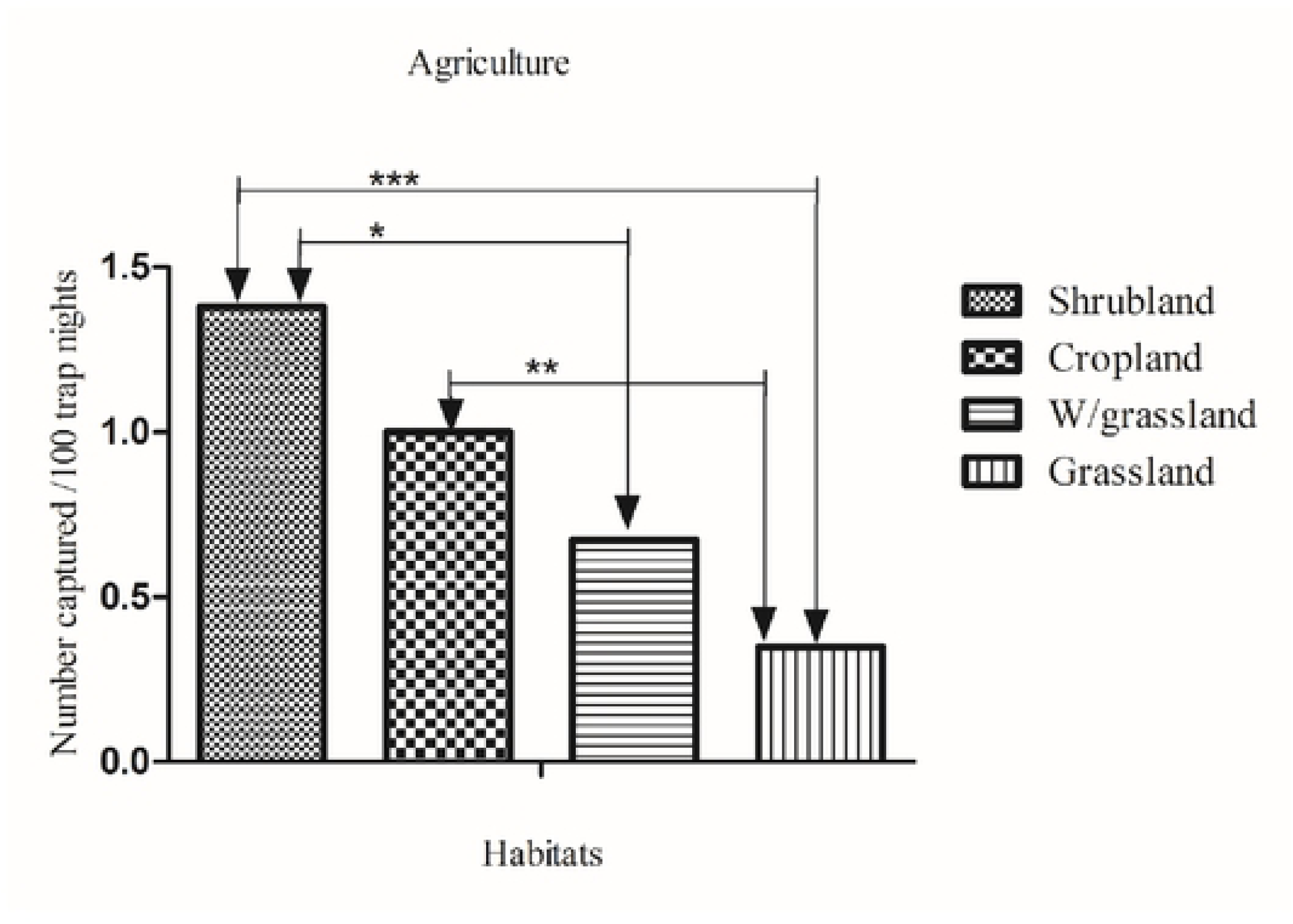
Number of small mammals caught per 100 trap nights (abundance) in each habitat in the AG landscape. Habitats with different numbers of small mammals are connected with a bar and an arrow. * Indicates level of significance (*P<0.01, **P<0.01, ***P<0.001).

The grassland habitat had a higher abundance of small mammals in the NP than the AG landscape 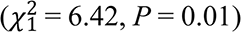. Similarly, small mammal abundance in the shrubland habitats varied across the three landscapes 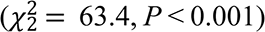 such that it was higher in the PA than the AG 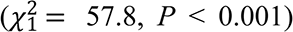 or the NP landscape 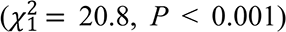. Also, the abundance of small mammals was higher in the shrubland habitat in the NP than in the AG landscape 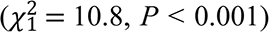. Although the shrubland habitat in the PA had the highest abundance of small mammals, it had relatively fewer species, with a single species dominating abundance in the habitat (78% of the total captures were of a single species), than it did in the NP or AG landscapes (Table A2). For the cropland habitat, the abundance of small mammals was higher in the PA than the AG landscape 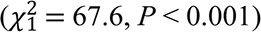. Likewise, abundance was higher in the wooded grassland habitat in the NP than the AG landscape 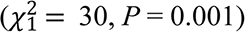

Small mammal abundance also varied interannually and seasonally (Fig. 7). Across all species, abundance was higher in 2018 than 2017 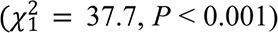 and in the dry than the wet season across both 2017 and 2018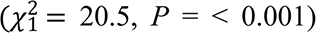. The seasonal variation in abundance persisted even when the two years were considered separately such that abundance was higher in the dry than the wet season in both 2017 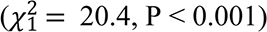 and 2018 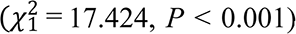. The number of individual species also varied seasonally with contrasts apparent across species. Collectively, *A. niloticus* 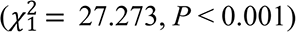, *M. natalensis* 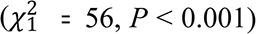 and *G. vicinus* 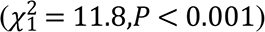 were more abundant in the dry than the wet season. By contrast *G. murinus* 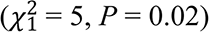 and *Crocidura spp* 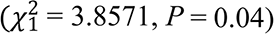 were more abundant in the wet than the dry season. All the other species (*Saccostomus sp, O. angoniensis, S. parvus, Zelomys sp, Zelotomys sp and P. jacksoni)* had lower capture rates and were mostly captured in the dry season in 2018.

**Fig 7.**
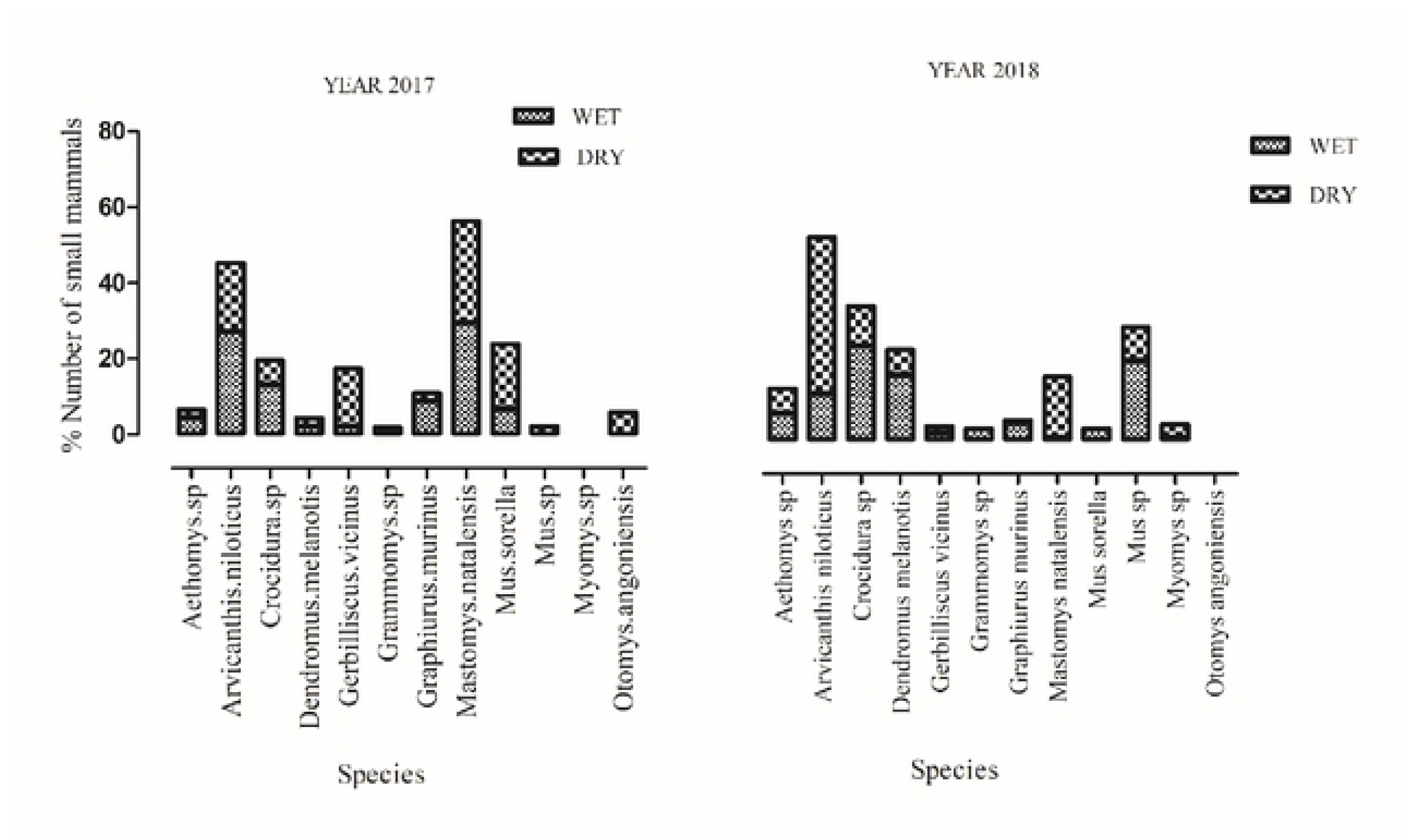
Percentage of all the small mammals caught in each season during 2017-2018 in the Serengeti ecosystem. Percentages are used here because the total number of trap nights was the same for both seasons.

## Discussion

### Small mammal species richness and diversity

As expected, species richness and diversity of small mammals were higher inside the NP than in either the AG or PA landscape. The higher diversity in the NP demonstrates that protection is crucial in safeguarding wildlife. This is further reinforced by the observation that most of the species that had low trap success occurred in the NP, indicating speciality. Furthermore, the NP is the least modified by human activities and thus has high vegetation heterogeneity and intactness, crucial to supporting a variety of small mammal species. Habitat heterogeneity is one of the most important factors influencing small mammal richness and diversity (50, 51). These findings concur with those of Magige and Senzota (2006) who also recorded the highest small mammal diversity in the protected landscape. They are also consistent with the general notion that greater habitat diversity is associated with higher species diversity (52)

Similarly, the higher species diversity for the NP landscape reflects the importance of protection in maintaining high habitat and species diversity in ecosystems. High habitat diversity is essential to high species diversity because the presence of habitat specialists is conditional on the presence of their favoured habitat types. Thus, for example, the riverine forest habitat, harbored mainly *G. murinus* and *Grammomys spp*, both of which prefer trees and intact forest cover for nesting in the Serengeti. The selection of the riverine forest habitat by *G. murinus* has also been noted previously (53) and is indicative of habitat specificity. The impact of livestock grazing in the shrubland habitat in the PA landscape was manifested in the lower small mammal species diversity than in the other habitats. Moreover, the relatively lower evenness (30%) in the PA landscape reaffirms the role of livestock grazing as one of the anthropogenic activities that alter vegetation structure and promote generalist species over habitat specialists. So, how does livestock grazing reduce small mammal species diversity? One plausible mechanism is that grazing increases shrub cover and patches and hence nesting and refuge sites for small mammals but reduces vegetation diversity and ground cover (54). Thus, continuous grazing decreases small mammal species diversity by reducing their food diversity and increasing predation risk (29, 55). In consequence, human activities, such as livestock grazing, are detrimental to ecosystems as they reduce small mammal species diversity, yet small mammals play a central role in food webs and other ecosystem services.

### Small mammal abundance and species composition

The fact that small mammal abundance was the highest in the pastoral, middling in the protected and the lowest in the agricultural landscape deviates from our initial expectation that abundance should be the highest in the park, intermediate in pastoral and the least in the agricultural landscape. This deviation is primarily attributable to the dominance of *A. niloticus* and *M. natalesis* in the pastoral landscape. When either one or both species (*A. niloticus and M. natalensis*) are excluded from analysis, then the results conform to our prediction, implying that the two numerically dominant species make the pastoral landscape to have more abundant but fewer species. The latter two species are generalists able to produce many young, attain high densities in relatively short time frames and colonize new areas (56, 57). The numerical dominance of these species suggests that human activities might have modified habitats in pastoral lands, rendering them suitable for a few generalist species (14, 17).

Small mammal abundance also varied across habitats and was the highest in the shrubland habitat in the pastoral landscape. Notably, *A*. *niloticus* (78%) was the most abundant species in the shrubland habitat in the pastoral landscape where sustained livestock grazing may have resulted in increased woody plant (shrub) cover (58, 59) at the expense of herbaceous plant cover. This accords with the observation that heavy livestock grazing can reduce plant species diversity and homogenize natural habitats (29, 54, 60). By increasing shrub cover and reducing primary productivity of the above-ground biomass, livestock grazing can negatively affect plant diversity and food availability for small mammals. On the other hand, small mammals were also abundant in the cropland habitat in the pastoral landscape due primarily to the numerical dominance of *M. natalensis* in the post-harvest (dry season) period when food and cover are still relatively plentiful. This species is common in croplands due to its feeding ecology, generalist behaviour and high food availability in this habitat after harvests (18, 27, 56, 57).

In contrast, the cropland habitat had significantly lower overall abundance in the agricultural than the pastoral landscape due primarily to differences in the cropping systems used in the two landscapes. In the pastoral landscape, crop farms are fenced to prevent livestock from raiding crops. The vegetation fringing the fences provide habitats for small mammals. In addition, during harvests, crop farmers in the pastoral landscape often leave some maize stocks and cobs on the farm for livestock. Also, after harvests farmers in the pastoral landscape take relatively longer time before preparing land by hand hoes for the next planting season. Thus, the land remains relatively intact for some months. By contrast, fencing farms is not common in the agricultural landscape, and it typically takes less than a month to prepare land by oxen and replant because most of the farmers cultivate crops for cash income (Pers. Obs &comm. 2018). Thus, the cropping system used in the cropland habitat in the pastoral landscape likely contributed to a relatively stable supply and availability of food and shelter for the small mammal species after harvests. Preparing land and replanting soon after harvesting, as done in the cropland in the agricultural landscape, reduces shelter and likely exposes small mammals to high predation risk, forcing them to seek safer habitats elsewhere. Changes in the quality and quantity of resources associated with cultivation can thus greatly influence the population size of small mammals. Specifically, the land preparation methods used in the cropland habitat in the pastoral landscape may favour *M. natalensis* species because it disturbs the habitat much less than a tractor or oxen does (61).

The higher abundance of small mammals in both the wooded grassland and grassland habitats in the NP than the AG landscape may reflect greater habitat heterogeneity in the NP landscape because of less human activities (17). Specifically, the higher abundance of *D. melanotis* in the wooded grassland and shrubland habitats in the NP than in the AG landscape could be due to advanced locomotor adaptations and mode of foraging of *D. melanotis.* This species is an agile climber that prefers tall vegetation (62) prevalent in the NP landscape. Although it occurs in both relatively pristine and human-dominated habitats, disturbance such as from agriculture, may cause their temporal migration (63). Thus, the NP is likely safer than the human-dominated landscapes besides providing tall vegetation habitats favoured by this species.

The higher abundance of small mammals in the dry than the wet season is consistent with the expectation. Seasonality alters vegetation cover and food availability and thus small mammal abundance (17, 57). Food and shelter rank among the key factors that determine small mammals abundance, hence the higher abundance in the dry season indicates elevated food availability due to high rainfall in the preceding wet season (64, 65). Surprisingly, *Crocidura spp* and *G. murinus* were more abundant in the wet than the dry season. This contradicts findings of two previous studies which reported higher abundances of both species in the dry than the wet season in this ecosystem (66, 67), implying substantial interannual variation in abundance. Such interannual fluctuations in abundance may be linked to a similar underlying variation in rainfall and hence in food availability.

In aggregate, these results support the notion that human activities, such as grazing and agriculture, homogenize habitats. This is demonstrated by the higher abundance of *A. niloticus* in the shrubland and *M. natalensis* in the cropland habitat. Both species are habitat generalists able to expand their home ranges depending on seasonal food availability and to persist in disturbed areas (57, 68, 69). This conforms with the general view that human-dominated habitats should harbour many generalist small mammal species (Byrom et al., 2015). Thus, by creating habitats that favor generalists at the expense of specialist species, human activities modify ecosystem function and suitability for small mammal communities.

### Conclusions and conservation implications

Human activities apparently exert deleterious effects on small mammals through reducing habitat suitability, resulting in reduced abundance, species richness and diversity. The protected area had higher species richness and diversity than the adjoining agricultural or pastoral landscapes, implying better protection of small mammals. Agricultural landscapes support less abundant but more diverse small mammal communities than pastoral landscapes due to greater food and shelter availability post-harvest than in the pastoral lands. Pastoral lands had more abundant but less diverse small mammal communities. This implicates structural habitat modification and loss of habitat heterogeneity. Loss of habitat heterogeneity due to human activities is associated with the loss of important habitats for small mammal species and hence with the loss of many small mammal species and the ecological services they provide. These findings reaffirm the importance of protection as a strategy for conserving the abundance, richness and diversity of small mammal species and can aid conservationists in diagnosing healthy ecosystems.

## Acknowledgements

We thank the Tanzania National Parks (TANAPA) and the Tanzania Wildlife Research Institute (TAWIRI) for permission to conduct this study.

## Supporting information

**S1 Table. Small mammal species richness and diversity and the significance of tests of their differences between habitats in three land use types in the Serengeti Ecosystem between 2017 and 2018. AG = Agriculture, NP = National Park and PA Pastoral landscape**

H’= Species diversity, S = Species richness, ns = non-significant, n = number of samples

* Indicates level of significance **P<0.01, ***P<0.001).

**S2 Table. Relative abundance (number of individuals of a species caught in a habitat divided by total number of individuals of all species caught in the habitat × 100) of small mammal species captured in 10 different habitats in the Serengeti ecosystem, Tanzania, during the wet and dry seasons of 2017 and 2018**.

^†^AG = Agriculture, NP = National Park, PA = Pastoral Landscape, Crop = cropland, Sh = Shrubland, WG = Wooded grassland, Gr= Grassland and For = Riverine forest.

